# Comparative Cilia Analysis in the Cerebral Cortices of Turtles, Mice, and Macaques

**DOI:** 10.1101/2025.11.16.688725

**Authors:** Soheila Mirhosseiniardakani, Liyan Qiu, Konstantinos Sousounis, Mark Lyon, Xuanmao Chen

**Affiliations:** Department of Molecular, Cellular, and Biomedical Sciences; College of Life Sciences and Agriculture, University of New Hampshire, Durham, NH 03824, USA; Department of Mathematics and Statistics, College of Engineering and Physical Sciences, University of New Hampshire, Durham, NH 03824, USA

**Keywords:** Primary Cilia, Principal Neuron Positioning, Cortical Evolution, Cilia Orientation, inside-out

## Abstract

Primary cilia are centriole-derived sensory organelles found in most vertebrate cells including neurons. In the mouse neocortex, the primary cilia of pyramidal neurons are found to orient predominantly toward the pia, reflecting a reverse movement that occurs during postnatal neuronal repositioning. This study compared cilia orientation in principal excitatory neurons in the cerebral cortex across turtles, mice, and macaques to identify patterns to infer mechanisms of cortical evolution. We first developed custom MATLAB Apps to facilitate the fast identification and statistical analyses of cilia orientation. We found that generally the primary cilia of principal neurons in sparse inside-out laminated regions, including the macaque and mouse neocortex, macaque CA1, mouse entorhinal cortex and neighboring regions, and mouse piriform cortex layer III, orient toward the pia. In contrast, primary cilia in compact laminae of these species, including the macaque and mouse dentate gyrus (DG), macaque CA3, mouse piriform cortex layer II, and turtle lateral cortex manifest opposite orientations, positioning perpendicular to the laminae. These data suggest that over the course of cortical evolution, primary cilia in principal neurons evolve from initially having no preferred orientation to becoming increasingly oriented toward the pial surface. We propose a working model for cortical evolution: the placement of principal neurons progresses from minimal migration in lower vertebrates to forward migration in higher species, and ultimately to pronounced reverse soma movement in higher inside-out laminated cortices, driven by the accumulation of large neuronal populations in the outer layers after completing radial migration.

## Introduction

The mammalian cerebral cortex are multilayered, radially organized brain structures, whose formation relies on the proper migration of principal neurons from the germinal zones to their final destinations^1, 2^. It encompasses the 6-layered neocortex, the 3- to 4-layered allocortices such as olfactory cortex and hippocampus, and neighboring structures ^2–7^. The sparsely layered neocortex is developed in an inside-out pattern^8, 9^. Similarly, the piriform cortex, a major part of the olfactory cortex, is also an inside-out laminated structure with a portion of principal neurons condensed into the compact layer II, while the rest distributed to the sparse layer III ^6, 10–13^. However, in the mouse hippocampus, the CA1 region has three layers, where almost all principal neurons are compacted into layer II (stratum pyramidal) in an inside-out manner ^14, 15^. Intriguingly, inside-out layered cortical structures vary widely in neuron density among the mammalian brains. In contrast, the outside-in laminated DG is highly conserved across mammals, with neurons densely packed in the principal cell layer ^16^.

Only mammals possess the sparsely laminated neocortex, which enables them to receive, process, and store a vast amount of information and perform higher-order brain functions ^6, 17^. The 6-layered neocortex in the mammalian brains evolutionarily correspond to the dorsal cortex in reptiles ^18, 19^. The 3-layered olfactory cortex and hippocampus in mammals are equivalent to the lateral and medial cortex in reptiles, respectively ^7, 18, 19^. The dorsal, ventral, and medial cortices of retiles thereby provide a valuable comparative model for studying the evolution of cortical lamination.

Primary cilia are centriole-derived sensory organelles that exist in most cell types of mammals, including neurons in the brain ^20–26^. Cilia are anchored to the cell at their base by the basal body and composed of a ring of 9 triplets of gamma-tubulin ^27^. The axoneme of cilia is composed of microtubules arranged in a radial array of nine doublets ^28^. Primary cilia detect various extracellular signals, including neurotransmitters, morphogens and hormones, and control a wide range of cellular functions including energy balance, neurogenesis and dorsal-ventral neural tube patterning^23, 25, 29–33^. Alterations in ciliary structure or signaling have been linked to neurodevelopmental abnormalities and are implicated in a subset of ciliopathies that affect brain development ^22, 23, 25, 34–36^. However, the function and acting mechanisms of neuronal primary cilia in the postnatal brain remain elusive.

We have previously reported that the primary cilia of pyramidal neurons in the deep and superficial sublayers in the mouse hippocampal CA1 region exhibit opposite orientation at P10, whereas in the neocortex, they predominantly orient toward the pial surface ^16^. The directional cilia orientation in inside-out laminated regions reflects reverse movement of principal neurons during early postnatal neurodevelopment ^16^. The reverse movement is considered crucial for the evolutionary transition from the compact, 3-layered allocortex to the sparse, 6-layered neocortex^16^. This study sought to determine cilia orientation patterns of principal neurons across the cerebral cortices of turtles, mice, and macaques to evaluate shared patterns and infer the mechanisms of cortical evolution.

## Materials and Methods

### Animals

The Arl13b-mCherry; Centrin2-GFP double transgenic mice (Stock No. 027967) ^37–39^ were obtained from the Jackson Laboratory (JAX) and originally carried C57Bl/6, BALB/c, and FVB/N mixed genetic background. As described previously, the expression of Arl13b-mCherry and Centrin2-GFP are both driven by the pCAGGs promoter, which contains a chicken *β*-actin promoter and CMV immediate early enhancer. After importing to the Chen laboratory, the line was backcrossed with C57BL/6J mice for over six generations to achieve a uniform C57BL/6J genetic background. This process produced three distinct lines: a double transgenic line, which expresses both Arl13b-mCherry and Centrin2-GFP and was used to visualize the orientation of primary cilia, as well as two single transgenic lines expressing either Arl13b-mCherry or Centrin2-GFP individually. Both male and female mice were included in the experiments. Mice were maintained on a 12-hour light/dark cycle at 22 °C and had access to food and water ad libitum. All animal procedures were approved by the Institutional Animal Care and Use Committee at the University of New Hampshire (UNH). 6 fixed adult turtle brain tissues from *Chrysemys picta bellii* were sent from the Gilles Laurent’s laboratory in Max Plank Institute for Brain Research in Germany. They were male and around 6 - 7 years old ^18, 19^. 9 coronal sections of adult macaque fixed brain tissues from 3 animals (mix sexes) were from Dr. Carol Barnes at the University of Arizona ^40–42^. All brain tissues, including the cortices and hippocampi, and their thickness was 30 μm. They were fixed in 4% paraformaldehyde (PFA) and 0.1 M PBS.

### Brain Tissue Collection

Mice were deeply anesthetized with an intraperitoneal injection of ketamine and xylazine and then were transcardially perfused with ice-cold 0.1 M phosphate buffer saline (PBS) followed by 4% paraformaldehyde (PFA) in 0.1 M PBS ^43^. Mouse brains were extracted and then placed into 4% PFA overnight at 4 °C. The following day, mouse and turtle brains were rinsed in PBS for 30 min at room temperature. Mouse and turtle brains then were dehydrated with 30% sucrose in 0.1 M PBS overnight at 4°C. Finally, those tissues were embedded in O.C.T. resin before being sectioned at −18 °C using a cryostat to a thickness of 50 μm. Before staining the macaque brain tissues, an antigen retrieval step was used in most formalin-fixed tissues to reverse epitope masking and restore epitope-antibody binding often lost during the fixation process.

### Immunofluorescence Staining

Mouse and turtle brains sections were sliced and placed in 24-wells plate, staying afloat during the staining procedure ^16^. Macaque brain tissues were cut into 4 sections evenly as they were bigger than mice and turtles’ tissues and placed in a 6-wells plate. All tissues were washed three times with 0.1 M PBS. Then they were washed in 0.2% Triton X-100 (vol/vol) in 0.1 M PBS three times for 10 minutes at room temperature. Then they were blocked in blocking buffer (0.1M PBS containing 10% normal donkey serum (vol/vol), 2% bovine serum albumin (weight/vol), and 0.2% Triton X-100 (vol/vol)) for one hour at room temperature. After that, slices were incubated with primary antibody overnight at 4 °C in a blocking buffer. After washing with 0.1 M PBS three times for 10 minutes and then slices were incubated with secondary antibodies for 1 h at room temperature. Cilia can be marked by antibodies of cilia-specific proteins, including adenylyl cyclase subtype 3 (AC3). Primary antibodies were rabbit anti-AC3 (1:10000, EnCor Biotechnology Inc., #RPCA-ACIII), and anti-NeuN (EMD Millipore Corp., #3612227). NeuN antibody marks mature cells. Secondary antibodies were Alexa Fluor 488-, 546-, or 647-conjugated (Invitrogen, La Jolla, CA). Finally, sections were counter-stained with DAPI (Southern Biotech™, #OB010020) and images were acquired with the confocal microscope (Nikon, A1R-HD).

### Cilia Identification and Orientation Measurement

A custom MATLAB® Apps, named Cilia Locator, was created to facilitate the fast identification of primary cilia in multiple layers of confocal images and to determine their orientation. Details of the new method are described in Results Section and Fig. 1A.

**Fig. 1:**
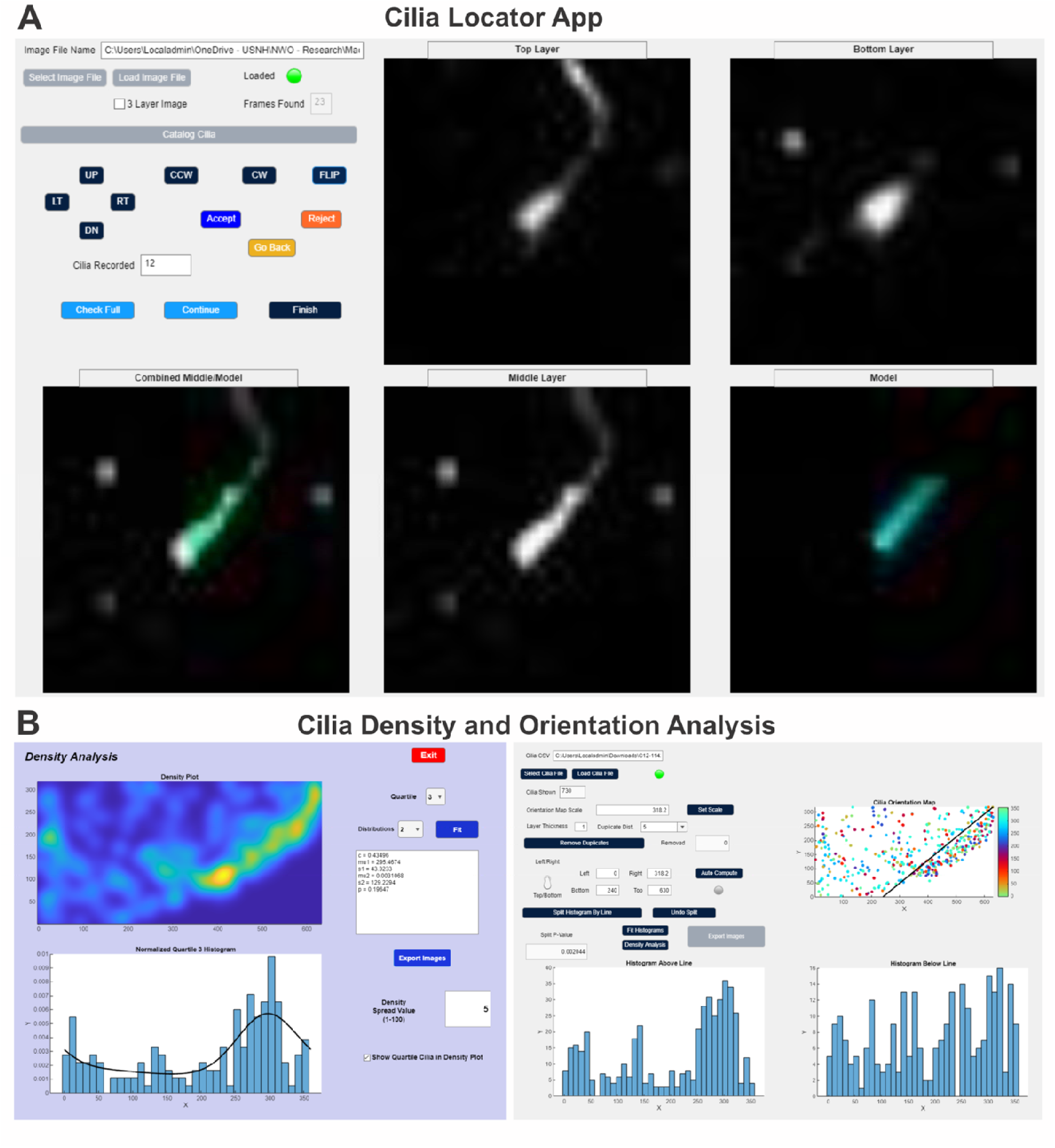
MATLAB Apps of cilia identification and statistical analysis. **(A)** Cilia Locator App automatically identifying and orienting cilia from multilayer confocal images. **(B)** Cilia Analysis App demonstrating various analysis tools for the statistical analysis of cilia including correlating the orientation distribution to the density.

### Cilia Orientation Distribution Fitting

An additional custom MATLAB® App, named Cilia Analysis, was developed to analyze the catalogued cilia identification and orientation data. Cilia orientation distribution fitting method is described in Results Section

### Primary Cilia Length Measurement

The freehand line tool within Fiji (ImageJ) was applied to measure cilia length from the base of axoneme to the distal tip. 50-μm Z-stack confocal images were used for Arl13B-mCherry mice and turtles. 30-μm Z-stack confocal images are used for macaques.

### Cilia Frequency Measurement

Cilia frequency is calculated by cilia number divided by nuclei number in an image and then the same thing was done for 6 images^16^. An average of 6 numbers was measured and multiplied by 100 to present that in a percentage.

### Relative Cilia Diameter Estimation based on confocal images

Confocal images were zoomed in in Fiji (ImageJ) and the base of a single cilium, a thick part of the cilium, was selected. The freehand line tool within Fiji was used to measure cilia diameter from one side to the other side. Then, Ctrl and M bottoms were pressed at the same time. Consequently, one small window popped up and cilia measurements were shown. Those steps were done for 50 cilia, and the average of those cilia diameters was obtained for newts, turtles, macaques, and mice.

### Statistical analysis

Data were analyzed with Fiji (ImageJ), GraphPad Prism and custom MATLAB Apps. Data was considered as statistically significant if p < 0.05 and values in the graph are expressed as mean ± standard error of the mean (SEM). Statistical analyses were conducted using one-way ANOVA for multiple group comparison, as appropriate. If not otherwise indicated in the main text or figure legends, N.S., Not Significant; * p < 0.05; ** p < 0.01; *** p < 0.001.

## Results

### Automatic Cilia Identification and Orientation Measurement by *Cilia Locator*

A custom MATLAB® App, named *Cilia Locator*, was created to facilitate the fast identification of cilia and to determine their orientation. Acquired confocal images comprising multiple monochromatic layers were loaded into the application. Letting *B(i, j, k)* be the brightness of the pixel located at coordinates

*(i, j)* on layer *k*, the algorithm proceeds by using a model consisting of just few pixels (*NX M*) and layers (*Q*), based on the resolution of the image, for the cilia’s nucleus. We denote the model *H(n, m, q)* and then we may compute:

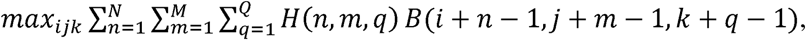

for values of *i, j,* and *k* sufficiently far from the borders of the image that cilia can reasonably be detected (Fig. 1A). The maximum product sum, which will occur when the model best correlates with the image, denotes the coordinates within the image of the most likely candidate for the nucleus of a cilia. An expanded cilia model for a full cilia, *E(n, m, q, θ)* oriented at an angle *θ* is then compared to the image at the site of the suspected nucleus by computing:

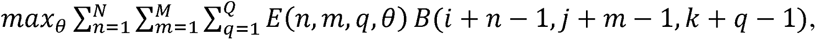

determining the most likely orientation of the full cilia. We note that the cilia nucleus model, *H(n, m, q)*, and the full cilia model, *E(n, m, q, θ)*, must be defined correctly in order to match the cilia at the magnification of the recorded image. As the magnification in this study was consistent, the same two models were used throughout.

After identifying a potential nucleus and orientation, the user of the software would be presented with several images including a view of the nucleus location on each of the three most relevant layers and an additional image of the pixelated cilia model overlayed on top of the center layer image of the nucleus. The software then presented the user with several options. The user can shift the position of the nucleus and/or orientation of the cilia in case it appears there was a better fit than the one the algorithm had identified. The user could then accept the cilia location and orientation or reject the location if it appeared the algorithm had misidentified some other aspect of the image as a nucleus. In either case, the results are catalogued, and that catalogue is also used to zero nearby values in *B(i, j, k)* so that the location (within at least two layers to either side of the center layer) will not be chosen again. At this point the algorithm would repeat with the computations as before, finding the next most likely nucleus candidate and determining the most likely orientation. When the user is next presented with images to confirm the location and orientation of the next cilia, previously identified cilia are marked in the images in a different color to avoid any double counting of cilia. The process continues until the algorithm is unable to find additional cilia and the user ends the identification process. The software also uses the catalogue to allow an undo command to go back in case a cilium was accepted or rejected in error (Fig. 1A).

Advantages of using the software is that it significantly sped up the identification as well as orientation measurement of the cilia. It also automatically zoomed in on the cilia during the cataloging process, minimizing the possibility of confirmation bias when different regions were expected to exhibit different orientations. Further, a larger number of cilia were able to be identified from each image with minimized risk in double counting of any cilia.

### Orientation Distribution Fitting by *Cilia Analysis*

An additional custom MATLAB App was developed to analyze the catalogued cilia identification and orientation data. The software utilized MATLAB’s custom distribution fitting algorithms to facilitate the fitting of the orientation data to several different distribution models. The Apps can fit cilia orientation data (0° - 360°) using one or two periodic normal distribution density functions, if there are possibly one or two major cilia orientations, or combined with a uniform distribution if a portion of the primary cilia do not have a specific orientation. The fitting will generate a peak value and standard deviation for one major orientation, and the percentage of cilia that point in that direction and evaluate the goodness of fitting by a Kolmogorov-Smirnov (K-S) test. The Apps also allow us to distinguish compact from sparse laminae (such as layer II vs. layer III of the mouse piriform cortex) based on cell/cilia density and separately calculate cilia orientation distribution in each layer. For each distribution fitting, the Kolmogorov–Smirnov test was applied to test the hypothesis that sampled orientation distribution could have been drawn from the fit probability distribution and facilitated the plotting of both the sampled orientation distribution and fit probability distribution (Fig. 1B). Additionally, the software included a feature to compute an average distribution density.

### Cilia orientation in various mouse cortical regions

We previously reported that the cilia orientation of principal neurons in the mouse cerebral cortex exhibits a certain pattern: primary cilia orient toward the pia in loosely layered region such as neocortex, exhibit opposite directions in compact cortical regions, such as hippocampus, or have no preferred orientation in excitatory neurons in nucleated regions, interneurons, or astrocytes ^16^. To assess whether these cilia orientation patterns are conserved across vertebrate species, we conducted a comparative cilia analysis using brain samples from turtles, mice, and macaques.

We first examined cilia orientation across different regions of the mouse cerebral cortex using immunofluorescence staining of mouse brain sections with AC3 antibodies. Confocal images were analyzed to quantify the orientation of neuronal primary cilia in each region. In the piriform cortex layer II (Fig. 2A), the histogram of cilia orientation angles (0° - 360°) displayed two distinct peaks approximately 180° apart, about 60% of cilia oriented toward the pial surface (∼270°), while ∼40% pointed in the opposite direction (∼90°). The corresponding polar plot confirmed this opposite orientation pattern. In contrast, the piriform cortex layer III (Fig. 2B), which has a sparsely laminated structure, showed a single dominant peak with ∼80% of cilia oriented toward the pia. The data fit a periodic normal distribution combined with a uniform component, and the polar plot confirmed a unidirectional orientation toward the pial surface. Similarly, in the cingulate cortex (Figs. 2C and 2D), both layers II and III displayed single-peaked histograms with most cilia concentrated around the major orientation. Approximately 60% of layer II and 90% of layer III cilia pointed toward the pial surface, as shown by the polar plots. Together, these findings indicate that cilia in the compact piriform layer II exhibit opposite directions while sparse cortical layers predominantly oriented toward the pial surface.

**Fig. 2.**
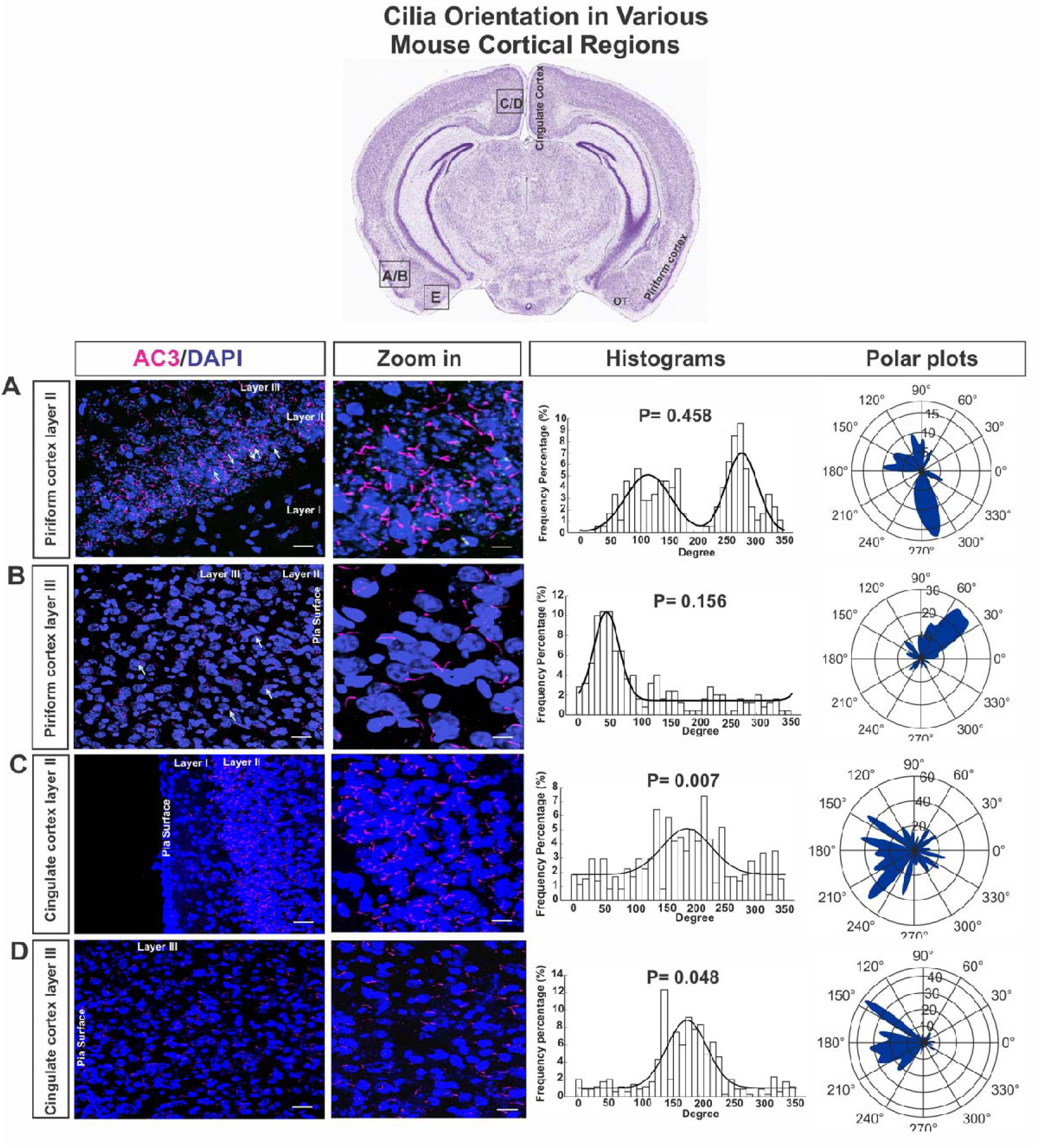
Cilia orientations in the piriform and cingulate cortex. At the top, a coronal section (adapted from the Allen Mouse Brain Atlas) shows the locations of examined cortical regions. (A) In layer II of the piriform cortex, neuronal cilia (AC3, pink; nuclei, blue) show mixed orientations, with peaks around 90° and 270° in the histogram (10° bins). The polar plot compares cilia oriented toward the pia surface (0 -180°) and the opposite direction (180 - 360°). (B) In layer III of the piriform cortex, most cilia point toward the pia, forming a single histogram peak; the polar plot confirms this predominant orientation. (C) In the cingulate cortex layer II, AC3-labeled cilia mainly point to the pia, as shown in both histogram and polar plot. (D) In layer III of the cingulate cortex, cilia orientations resemble layer II, with a single peak toward the pia. Scale bar = 50 μm.

We also examined cilia orientation in adjacent neocortical regions. In the entorhinal cortex (Figs. 3A, 3B), neuronal primary cilia in layers II/III and V showed a single peak in the orientation histograms, well fitted by a periodic normal distribution combined with a uniform component. Polar plots confirmed that most cilia were directed toward the pial surface. Similar results were observed in the perirhinal cortex (Figs. 3C, 3D), where ∼80% of cilia in layers II/III and V pointed to the pia. In the subiculum (Fig. 3E), which is a loose lamina, cilia orientation also displayed a single dominant peak (∼80% toward the pia). Together, these results indicate that primary cilia in both superficial and deep layers of the entorhinal, perirhinal, and subicular cortices are predominantly oriented toward the pial surface, consistent with the previous report ^16^.

**Fig. 3.**
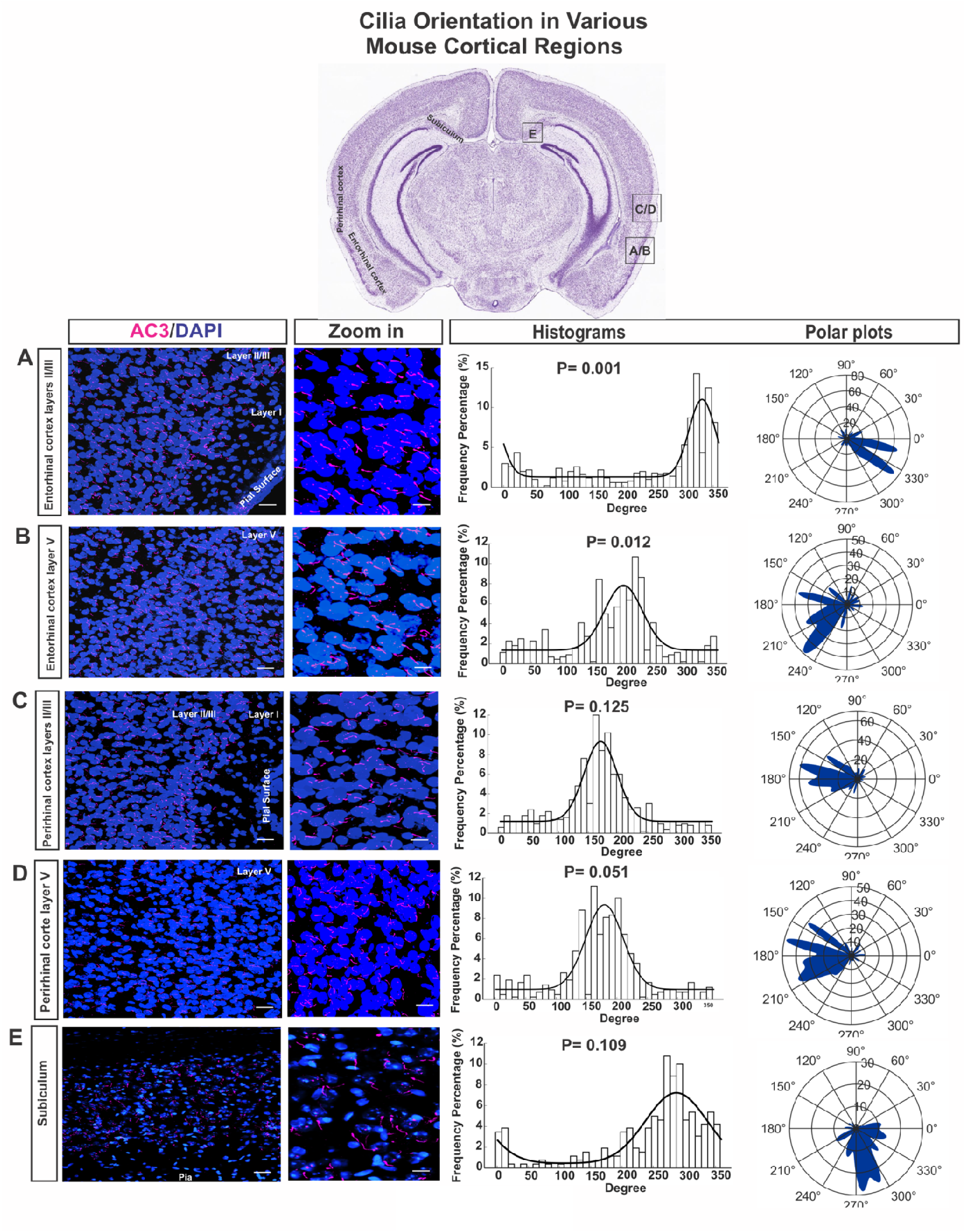
Neuronal primary cilia in the entorhinal and perirhinal cortices predominantly orient toward the pial surface. (A) Confocal image of neuronal cilia (AC3, pink; nuclei, blue) in superficial layers (II/III) of the entorhinal cortex shows most cilia pointing to the pia, confirmed by a single histogram peak and polar plot. (B) In the deep layer (V) of the entorhinal cortex, cilia exhibit a similar predominant orientation toward the pia. (C) In superficial layers (II/III) of the perirhinal cortex, cilia mostly point to the pia, as shown in the zoomed-in image, histogram, and polar plot. (D) In deep layer (V) of the perirhinal cortex, cilia also align toward the pia. (E) In the subiculum, most cilia show a single directional peak toward the pia surface. Scale bar = 50 μm.

### Cilia orientation in the turtle brain

We next analyzed cilia orientation across cortical regions in the turtle brain. Fig. 4 presents the results for the lateral, dorsal, medial, and ventral cortices, as well as the dorsal ventricular ridge (DVR). The top panel shows a coronal section of the turtle brain with the examined regions labeled. Different from mammals, reptile cortices only have compact, 3-layered allocortices, all outside-in layers ^10, 44^. Interestingly, their principal cell layers, except for the lateral cortex, do not migrate much away from the neurogenic zone (Fig. 4 top). Each panel in Fig. 4 presents immunofluorescence staining of neuronal primary cilia (AC3 staining) and matured neurons (NeuN staining), along with zoomed-in images, orientation histograms, and polar plots (left to right).

**Fig. 4.**
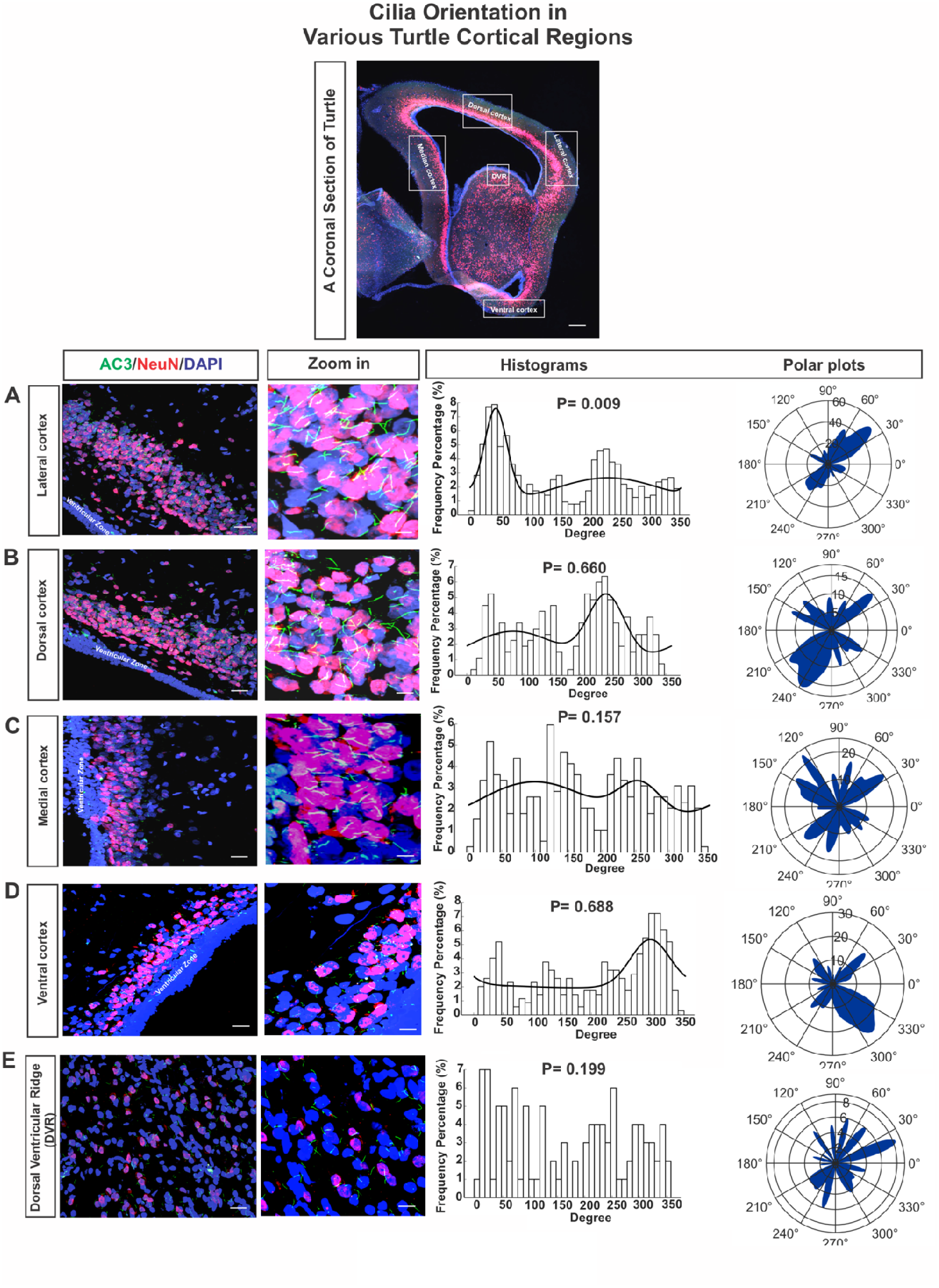
Cilia orientations in the turtle lateral, dorsal, medial, ventral cortices and dorsal ventricular ridge (DVR). At the top, a coronal section of the turtle brain shows the cortical regions analyzed. (A) In the lateral cortex, neuronal cilia (AC3, green; NeuN, red) show two main orientation peaks (∼50° and ∼220°), separated by ∼180°, as seen in the histogram (10° bins) and polar plot. The zoomed-in panel highlights orientations with some cilia pointing toward and others away from the ventricular zone. (B) In the dorsal cortex, cilia display a similar bidirectional pattern with orientations toward and away from the ventricular zone. (C) In the medial cortex, most cilia point toward the ventricular zone, forming a single dominant peak. (D) In the ventral cortex, cilia show two main orientations, partially directed toward the ventricular zone. (E) In DVR, cilia exhibit random orientations without a clear directional preference. Scale bar = 50 μm.

In the lateral cortex which localizes distantly from the ventricular zone (Fig. 4A and S1), the histogram shows two peaks roughly 180° apart, with ∼60% of cilia orientation concentrated around one peak and ∼40% around the other. The polar plot confirms that some cilia orient toward the ventricular zone, while others point in the opposite direction, indicating an opposite orientation pattern. In the dorsal cortex (Fig. 4B and S1), which do not migrate much, roughly two peaks, not sharp though, were observed (∼60° and ∼240°, around 180° apart), and the polar plot shows opposing cilia orientations - some directed toward the ventricular zone, others in the opposite direction. But the oppositie cilia orientation pattern was not strong (Fig. 4B and S1). In the medial cortex, which localized even more closely to the ventricular zone, primary cilia do not consistently align to one direction: some oriented toward the ventricular zone (Fig. 4C), others exhibit opposite direction, whereas the rest is parallel to the zone (Fig. 4C and S1). The orientation angle histogram was not well fitted by a mono or double periodic normal distribution density function. In the turtle ventral cortex (Fig. 4D and S1), the histogram of cilia orientation has one major peak, and one minor peak approximately 180° apart, which is not very strong. Finally, in the DVR (Fig. 4E), which is considered a non-laminated (non-layered) telencephalic pallial structure^44, 45^, cilia orientation appears irregular. The histogram shows no dominant peak, and the polar plot displays dispersed directions, indicating irregular cilia orientation. Together, these data show that turtle cortical regions display two opposite orientation patterns in the lateral cortex, somewhat opposite or irregular in the dorsal and median cortices, whereas the non-laminated DVR exhibits irregular orientation.

### Cilia in macaque cortices show opposite orientations in compact layers and a major orientation toward the pia in loose layers

We next examined cilia orientation in the macaque neocortex and hippocampus. The coronal section at the top of Fig. 5 labels the hippocampus and adjacent cortical areas. Panels display, from top to bottom, immunofluorescence images of neuronal primary cilia (AC3 staining), zoomed-in views, orientation histograms, and polar plots for the CA1, subiculum, CA3, and dentate gyrus (DG) regions. In the CA1 (Fig. 5A) and subiculum (Fig. 5B), histograms show a single dominant peak, with ∼90% of cilia clustered around it, well fitted by a single periodic normal distribution combined with a uniform component. Polar plots indicate that most cilia are oriented toward the pial surface. Notably, both CA1 and subiculum exhibit loose laminar organization, and their cilia orientations exhibit one major direction. In contrast, the CA3 (Fig. 5C) and DG (Fig. 5D) regions, which have denser laminar structures, show two distinct orientation peaks: ∼70% of cilia cluster around 90°, and ∼30% around 270° in CA3, while DG shows roughly equal distributions (∼50% each) at ∼90° and ∼270°, corresponding to directions approximately 180° apart. Polar plots confirm that a portion of cilia point toward the pia, while others orient in the opposite direction. Together, these findings reveal that cilia orientation in the macaque hippocampus differs by region: loosely layered CA1 and subiculum cilia predominantly align toward the pia, whereas compactly laminated CA3 and DG display opposing orientation patterns.

**Fig. 5.**
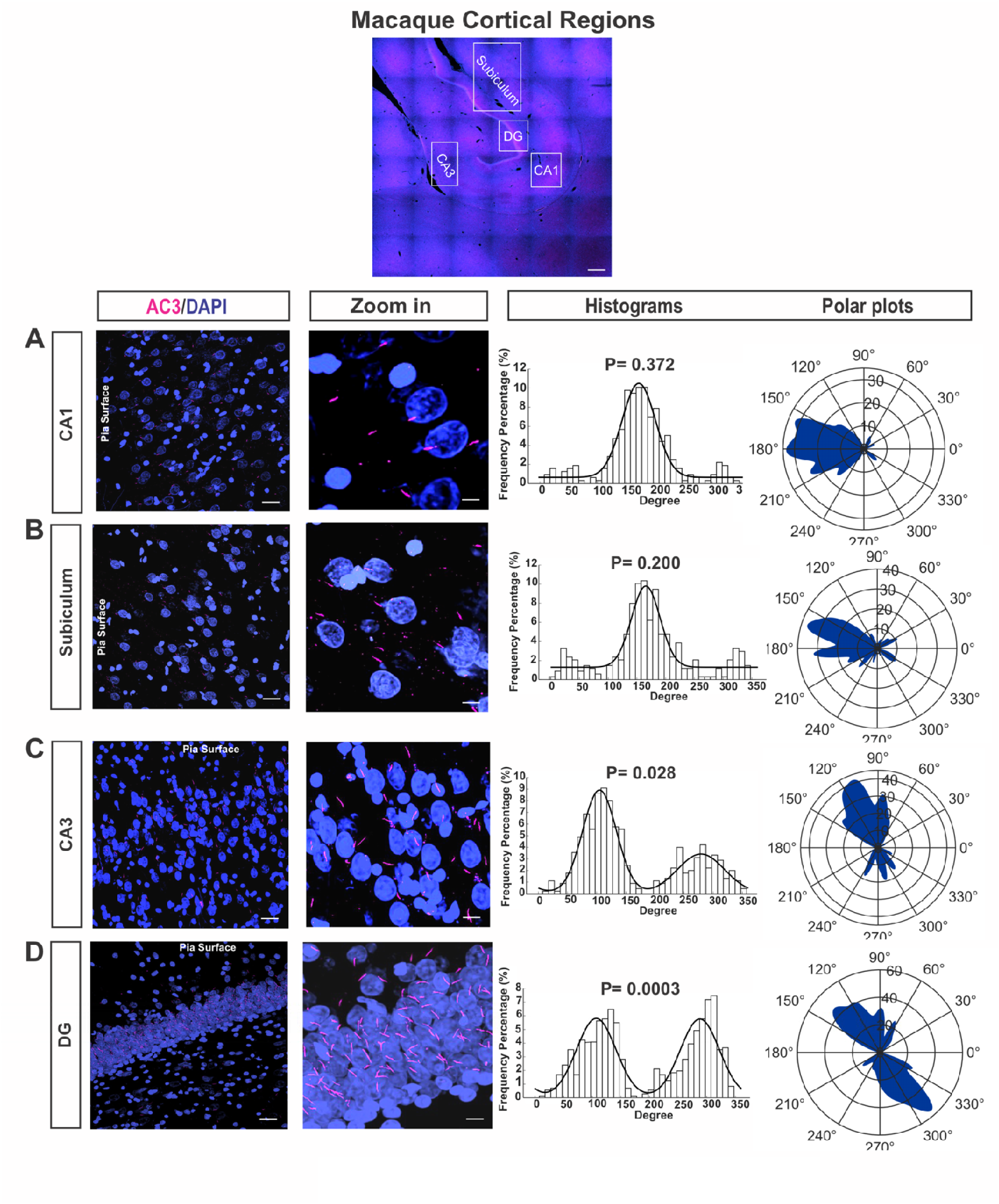
Cilia orientations in various macaque cortical regions. At the top, macaque brain sections show the hippocampus and cortical regions analyzed. (A) In the CA1 region, neuronal cilia (AC3, pink) mostly orient toward the pia surface, as shown in the zoomed-in panel, with a single histogram peak and a corresponding polar plot. (B) In the subiculum, cilia show a similar predominant orientation toward the pia, forming one clear peak in both histogram and polar plot. (C) In the CA3 region, cilia exhibit two main orientation peaks, indicating both pia-directed and opposite alignments. (D) In the dentate gyrus (DG), cilia also display two opposing peaks, with subsets pointing toward and away from the pia surface. Scale bar = 50 μm.

Fig. 6A and 6B further show data from the sulcal and gyral regions of the neocortex, both characterized by loose laminar organization. Each set includes immunofluorescence images (left), zoomed-in views of cilia, histograms of orientation angles, and corresponding polar plots (right). In the sulcus (Fig. 6A), the histogram displays a single peak with ∼90% of cilia clustered around it, and the polar plot confirms a predominant orientation toward the pial surface. Similarly, in the gyrus (Fig. 6B), the histogram fits a single periodic normal distribution combined with a uniform component, indicating that most cilia also point toward the pia. These results demonstrate that neuronal primary cilia in both sulcal and gyral regions generally orient toward the pial surface.

**Fig. 6.**
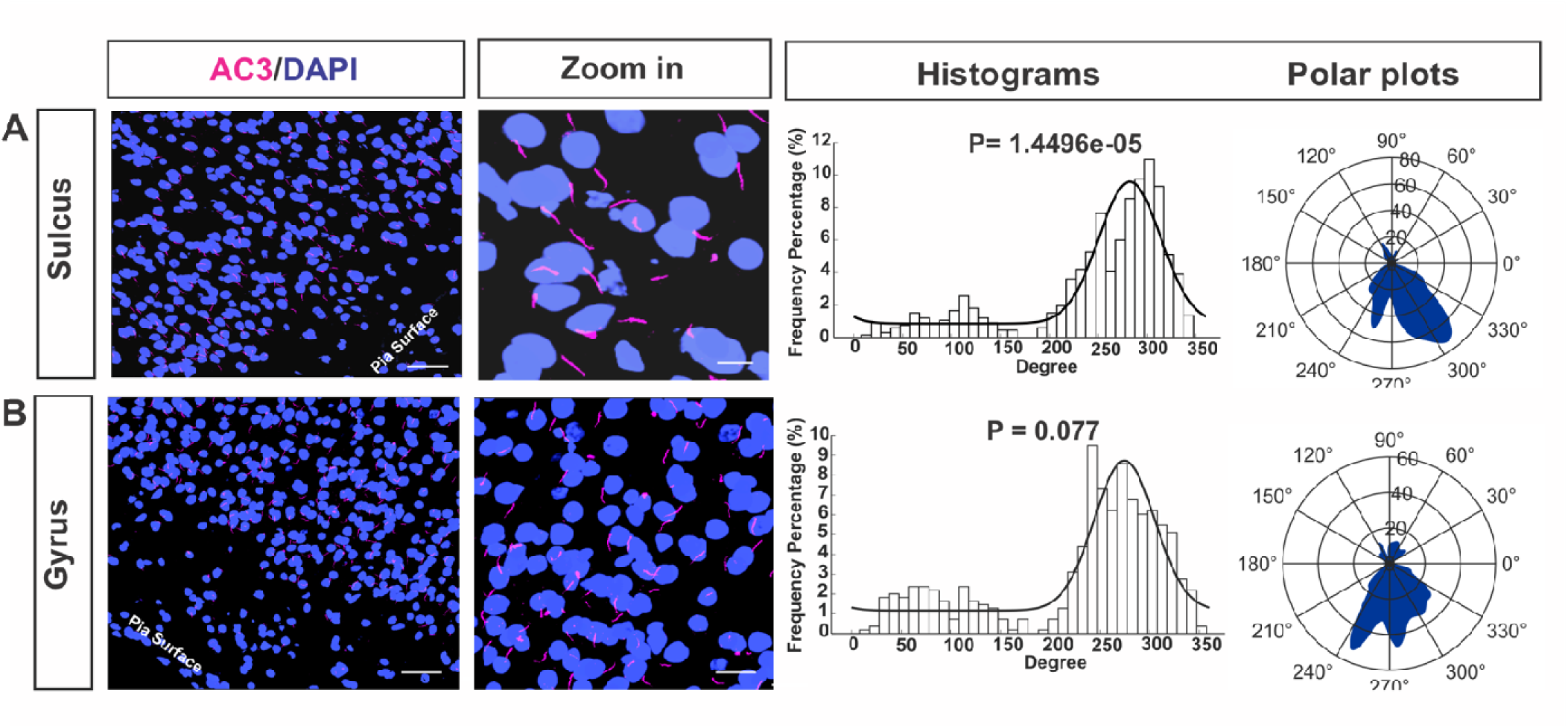
Cilia orientations in gyrified cortical regions of the macaque brain. At the top, orientation patterns of neuronal cilia across sulcal and gyral regions are shown, with most cilia pointing toward the pia surface. (A) In the sulcus, immunofluorescence staining of neuronal cilia (AC3, pink) reveals a predominant pia-directed orientation, as seen in the zoomed-in panels, histogram (10° bins) with a single peak, and polar plot. (B) In the gyrus, AC3-labeled cilia similarly show one dominant orientation toward the pia surface, confirmed by the histogram and polar plot. Scale bar = 50 μm.

### Cilia frequency is high in compact regions in turtle, mouse, and macaque brains

We also quantified cilia length and ciliation frequency in the turtle, mouse, and macaque brains. Fig. 7 summarizes these comparisons: the top bar graphs on the top show average cilia length, and the bottom bar graphs display ciliation frequency in various regions in turtle, mouse, and macaque brains. Ciliation frequency was highest in compact cortical areas: approximately 91% in the lateral cortex of turtles, 75% in the olfactory tubercle of mice, and 88% in the DG of macaques. These results suggest that compact cortical regions tend to exhibit higher cilia frequency across species.

**Fig. 7.**
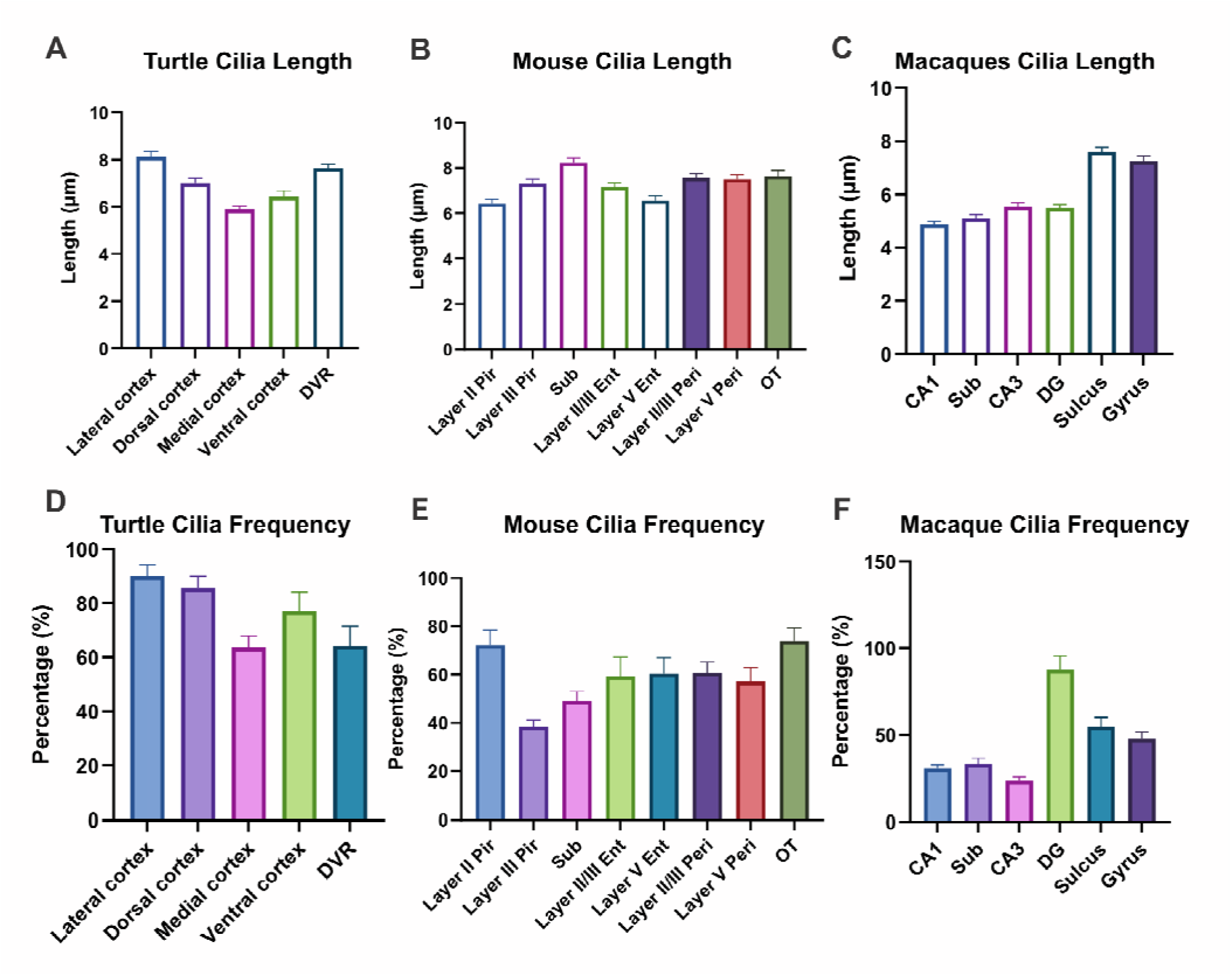
Cilia length and cilia frequency in different cortical regions of mice, macaques, and turtles. Bar graphs on the top exhibit cilia length in A) Mice, B) Macaques, and C) Turtles’ cortical regions. Bar graphs on the bottom show cilia frequency in A) Mice, B) Macaques, and C) Turtles’ cortical regions.

### Cilia diameter increases from reptiles to mammals

Relative cilia diameters were estimated from confocal images using Fiji (ImageJ). The average diameters were approximately 0.63 μm in turtles, 0.71 μm in mice, and 0.75 μm in macaques (Fig. 8). Among these species, macaques exhibited the thickest cilia, whereas tutles had the thinnest. Although confocal resolution limits precise diameter measurement, these data suggest a gradual increase in cilia diameter from reptiles to mammals.

**Fig. 8.**
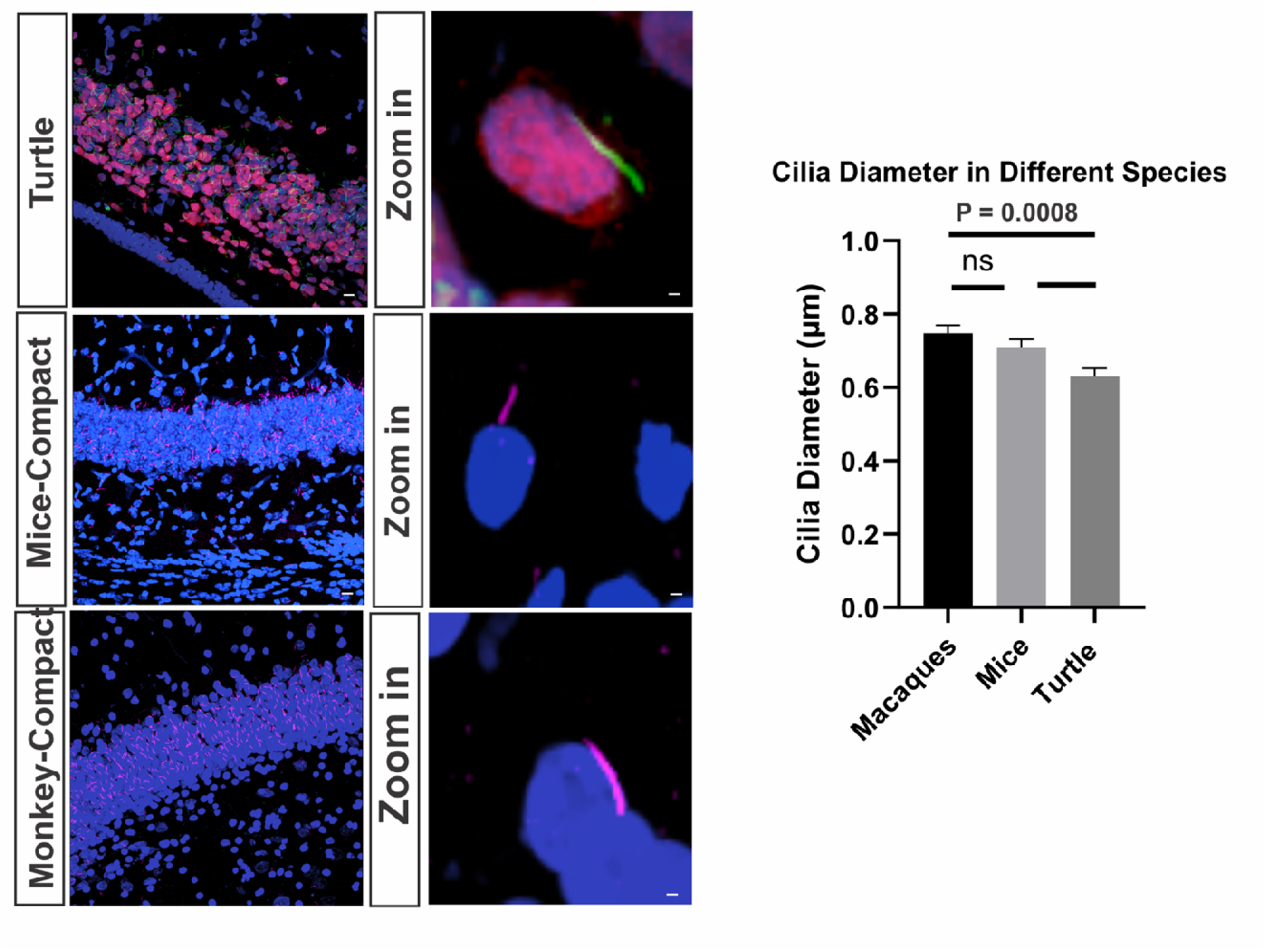
Relative neuronal cilia diameter in the turtle, mice, and macaque brains. Cilia diamete s of turtles, mice, and macaques were estimated based on confocal images. The exact diameters may be smaller.

### Arl13b and AC3 are co-expressed in neuroal primary cilia in the adult macaque brain

In the adult WT mouse brain, AC3 specifically labels neuronal primary cilia, whereas Arl13b marks astrocytic primary cilia, and the two proteins have minimal co-expression in neuronal and astrocytic cilia ^25, 38^. Interestingly, the expression patterns of AC3 and Arl13b in the adult macaque brain were different from mice. Fig. 9 shows that both AC3 and Arl13b are strongly expressed in neurons across multiple cortical regions, including CA1, CA3, DG, subiculum and neocortex. However, the enrichment of Arl13b in primary cilia is lower than that of AC3, and Arl13b is also distributed to non-ciliary subcelluar regions (Fig. 9). Arl13b is also expressed in astrocytic primary cilia. Fig. S2 presents images of Arl13b and GFAP immunostaining, demonstrating Arl13b expression in astrocyte primary cilia in different regions.

**Fig. 9.**
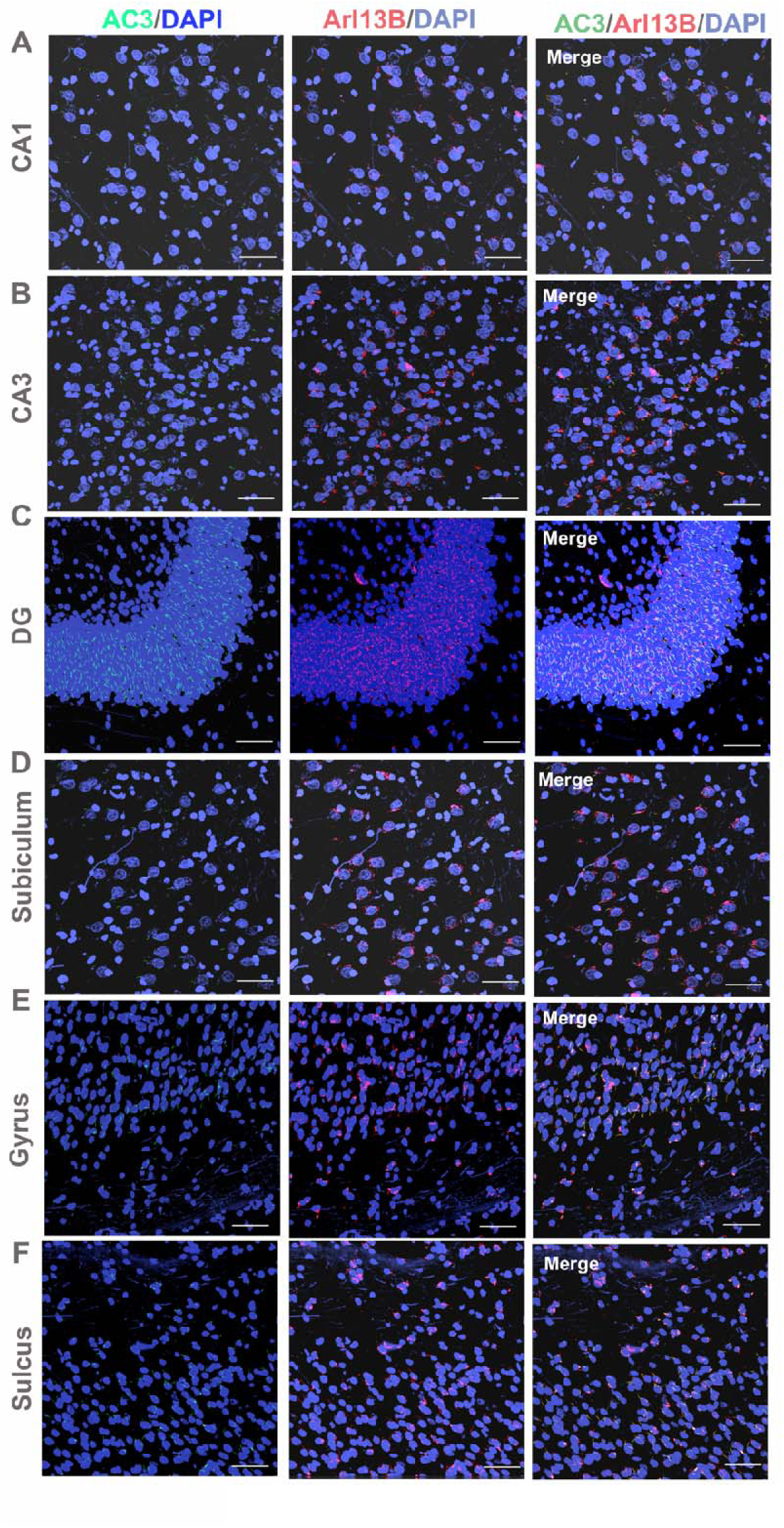
Both Arl13B and AC3 are expressed in neuronal primary cilia in the macaque brain. Arl13B and AC3 expressions in different regions of the adult macaque cerebral cortex. Arl13B is highly expressed in adult macaque neuronal primary cilia. The left panels are the merged images of AC3 and Arl13B Abs. They indicated that these two antibodies overlapped in all cortical regions. Scale bar = 50 μm.

## Discussion

Primary cilia are centriole-derived organelles found in most vertebrate cells, including neurons. While their basic structure are evolutionarily conserved across species ^46^, their specific roles and characteristics may differ. We compared cilia orientation in turtles, mice, and macaques and found that across species, compact laminae in the mammalian brain typically exhibit opposite cilia orientations, whereas loose layers show a predominant alignment toward the pial surface, and nucleated regions lack specific cilia orientations.

A technical limitation in cilia research is the lack of tools for fast, accurate and unbiased quantification of cilia orientation. Manual measurements are time-consuming, subjective, and prone to sampling bias, especially when cilia are variably oriented within thick confocal stacks. To overcome this limitation, we developed two complementary MATLAB-based applications, *Cilia Locator* and *Cilia Analysis*, to automate the detection and orientation measurement of primary cilia (Fig. 1). *Cilia Locator* scans multilayer confocal images to identify potential cilia based on fluorescence intensity and morphology, providing the user with visual confirmation of each detected cilium and its predicted orientation. This semi-automated approach enables high-throughput and accuracy while minimizing double counting or user bias. Once identified, cilia orientation data are exported to the *Cilia Analysis* for statistical modeling, which fit the angular distribution using one or two periodic normal functions, combined with a uniform component. This fit distinguishes unidirectional, bidirectional, and random orientation patterns and quantifies the dominant orientation angle, variance, and goodness of fit through a Kolmogorov-Smirnov test. Compared with manual methods, these tools markedly improve the reproducibility, accuracy, and speed of cilia orientation analysis.

In turtles, compact cortical regions such as the lateral cortex exhibit bidirectional cilia orientations (Fig. 4 and S1). Cilia in the dorsal and median cortices also tend to orient oppositely or irregularly, much less distinct than in the lateral cortex, this is likely because neurons in these regions do not migrate away from the neurogenic zone (Fig. 4 and S1). In contrast, the dorsal ventricular ridge (DVR), which lacks lamination, clearly shows irregular cilia orientation. In mice, opposite cilia orientations occur in compact regions such as the piriform layer II, dentate gyrus, and CA1, whereas sparse regions, including the entorhinal, perirhinal, and subicular cortices, show a preferential alignment toward the pia. Similarly, in macaques, compact regions (CA3 and DG) display clear opposing cilia orientations, while loose regions (CA1, subiculum, gyrus, and sulcus) predominantly exhibit pia-directed cilia (Fig. 5). Notably, although the hippocampal CA1 region in mice is compact with opposite cilia orientations at P10, the corresponding region in macaques is loosely organized and shows unidirectional alignment. Together, these results reveal a conserved relationship between laminar density and cilia directionality: dense laminae exhibit bidirectional cilia alignment, whereas sparse laminae consistently display pia-directed orientation, when principal cell layers migrate away from the neurogenic zone.

Mammals, reptiles, and birds evolved from a common amniote ancestor. The cortex in reptiles is a 3-layered structure, and only layer II is dense and contains principal neurons ^3, 18, 47, 48^. Different expansions and independent evolution of pallia lead to the brain’s diversity in mammals and reptiles, and these expansions were accompanied by the emergence of new neuronal cell types ^18^. Many turtles have ciliated olfactory receptor neurons in their nasal epithelium cavities that help them detect odorant reception ^49^. Research on primary cilia in turtles is limited. This study is the first to report the presence of neuronal primary cilia in distinct cortical regions of the turtle brain. We found cilia orientation of principal neurons are not well organized or oriented as in mammals (Fig. 4 and S1).

The cerebral cortices of mice and macaques comprise both compactly and loosely laminated structures ^3, 50^. In mice, the neocortex consists of six sparse layers, while layer II of the piriform cortex, which is part of the olfactory cortex, and the hippocampus exhibit compact laminar organization. All mouse hippocampal regions, including DG, CA1, CA2 and CA3, are compact, 3-layered laminae. The mouse piriform cortex layer III is loosely layered though. In contrast, the macaque hippocampus contains both compact and loose structures (Fig. 5). Specifically, the dentate gyrus (DG) is the only compact laminar region in the macaque hippocampus, whereas the CA1 and CA2 regions are loosely organized structures and CA3 somewhere in between. Additionally, the macaque cortex features sulci and gyri with loose architecture. The orientation of neuronal primary cilia differs between cortical regions: generally, in compact areas of mammalian brains, cilia exhibit opposing orientations, while in loose regions, they direct toward the pial surface ^16^.

We also observed that cilia length varies across cortical regions and species. The longest cilia are found in the sulcus of the macaque neocortex, the lateral cortex of turtles, and the CA1 region of mice (Table S1). Cilia diameter also varies among species. Primary cilia are immotile organelles with a typical 9+0 microtubule axoneme; however, neuronal cilia in the turtle brain, which have the thinnest diameters, may deviate from this pattern and contain fewer microtubule doublets (e.g., 7+0 or 6+0). This possibility remains to be validated by electron microscopy. It is important to note that studies on primary cilia across species are limited, and potential structural or functional change requires further validation.

The observed difference in ciliary markers supports ciliary function divergence between species. In the adult mouse brain, AC3 mostly marks neuronal primary cilia while Arl13b primarily labels astrocytic primary cilia ^25, 38^. They are barely co-expressed in adult neuronal primary cilia. In contrast, both proteins co-localize in neuronal primary cilia throughout the adult macaque brain (Fig. 8), suggesting an expanded or distinct role for Arl13b in primates and broader ciliary functionality in the primate brain.

In conclusion, from lower vertebrate reptiles to primates, cilia orientation in principal neurons evolves from irregular, bidirectional to predominant pia-directed alignment. These findings support a model in which, over the course of cortical evolution, primary cilia in principal neurons progress from initially irregular to increasingly oriented toward the pial surface. This reflects a trend in cortical evolution: principal neurons show little migration in lower vertebrates, then exhibit forward migration away from the neurogenic zone in higher species and ultimately display a stronger propensity for reverse soma movement in the inside-out laminated higher cortical layers, where larger neuronal populations accumulate in the outer layers after completing radial migration.

## Funding and Acknowledgments

This research was supported by National Institutes of Health Grants P20GM113131-7006, R15MH126317, and R15MH125305 to X.C.; UNH CoRE PRP awards and Cole Neuroscience and Behavioral Faculty Research Awards and a COLSA Community of Teaching and Research Scholar Award to X.C. and a UNH Summer TA Research Fellowship award (STAF) to S.H. We are grateful to the University Instrumentation Center at UNH for A1R HD confocal imaging service. We sincerely thank Drs. Carol Barnes and Gilles Laurent for providing turtle and macaque brain samples for cilia analysis.

---

**Table S1:**
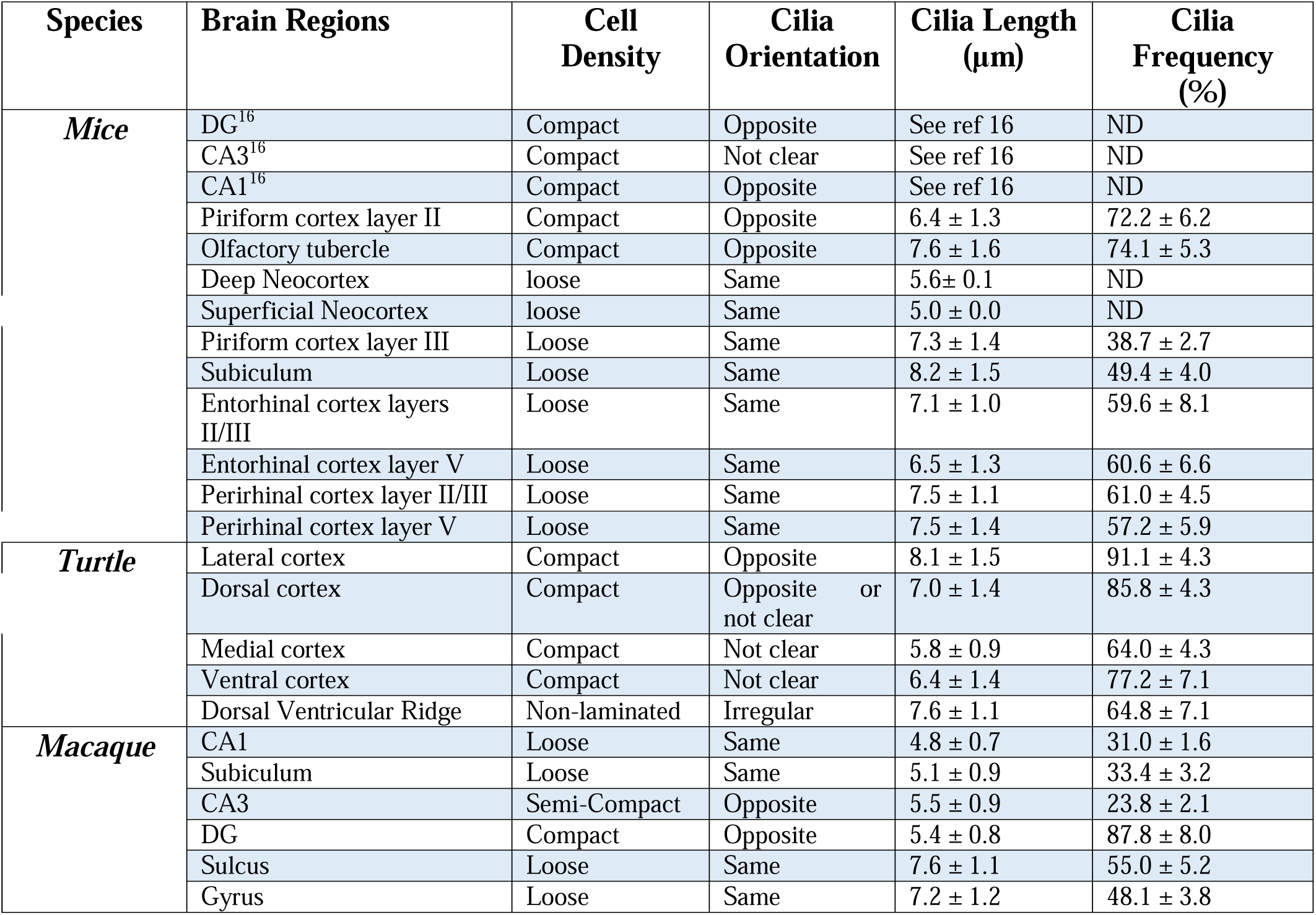

**Fig. S1.**
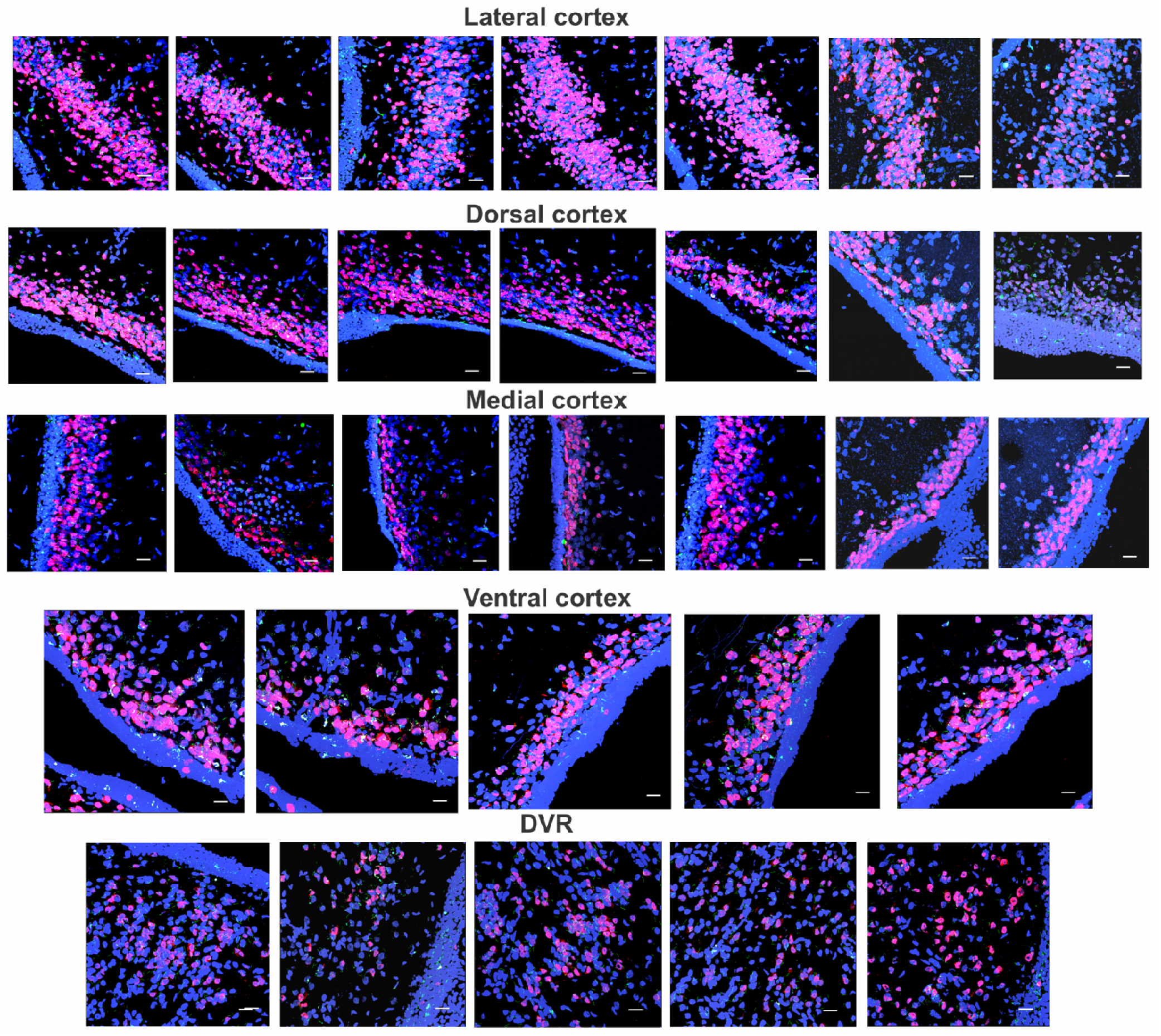
Examples of neuronal primary cilia images in different regions in the turtle brain, marked by AC3 and NeuN antibodies.

**Fig. S2.**
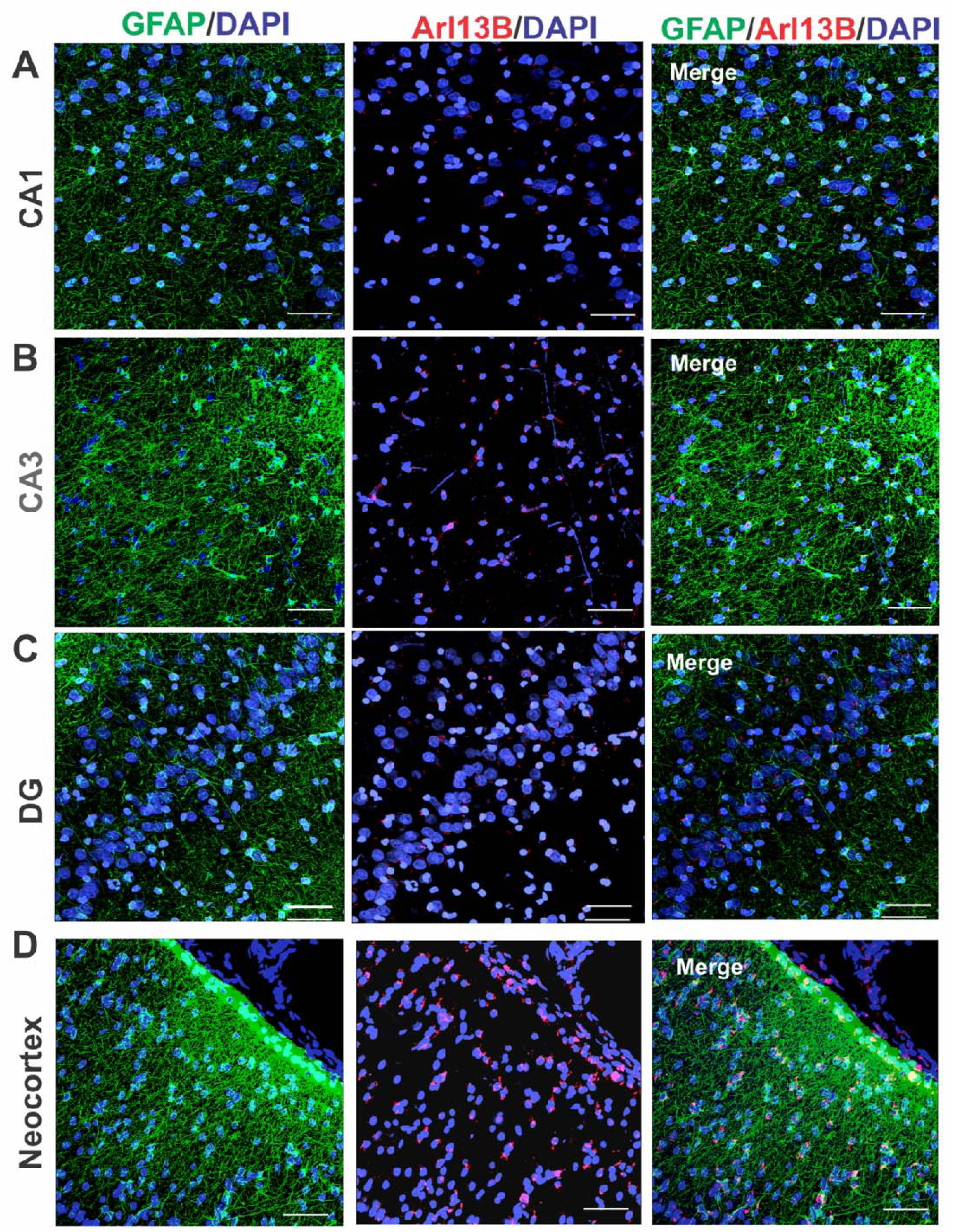
Arl13b is expressed in astrocytic primary cilia in the macaque brain. Arl13b and GFAP (Glial Fibrillary Acidic Protein) expressions are shown in different regions of the adult macaque cortex. Arl13b marked primary cilia and GFAP specifically labeled astrocytes. The right panels display merged images of GFAP, Arl13b, and DAPI. Scale bar = 50 μm. Clearly, GFAP-positive cells have Arl13b-marked primary cilia.

